# Denoising Approach Affects Diagnostic Differences in Brain Connectivity across the Alzheimer’s Disease Continuum

**DOI:** 10.1101/2022.06.16.496466

**Authors:** Jenna K. Blujus, Hwamee Oh, the Alzheimer’s Disease Neuroimaging Initiative

## Abstract

Graph theory provides a promising technique to investigate Alzheimer’s disease (AD)-related alterations in brain connectivity. However, discrepancies exist in the reported disruptions that occur to network topology across the AD continuum, which may be attributed to differences in the denoising approach used in fMRI processing to remove the effect of non-neuronal sources from signal. The current study aimed to determine if diagnostic differences in graph metrics were dependent on nuisance regression strategy. Sixty cognitively normal (CN), 60 MCI, and 40 AD matched for age, sex, and motion, were selected from the ADNI database for analysis. Resting state images were preprocessed using AFNI (v21.2.04) and 16 nuisance regression approaches were employed, which included the unique combination of four nuisance regressors (derivatives of the realignment parameters, motion censoring [euclidean norm > 0.3mm], outlier censoring [outlier fraction > .10], bandpass filtering [0.01 - 0.1 Hz]). Graph metrics representing network segregation (clustering coefficient, local efficiency, modularity), network integration (largest connected component, path length, local efficiency), and small-worldness (clustering coefficient/path length) were calculated. The results showed a significant interaction between diagnosis and nuisance approach on path length, such that diagnostic differences were only evident when motion derivatives and censoring of both motion and outlier volumes were applied. Further, regardless of the denoising approach, AD patients exhibited less segregated networks and lower small-worldness than CN and MCI. Finally, independent of diagnosis, denoising strategy significantly affected the magnitude of nearly all metrics (except local efficiency), such that models including bandpass filtering had higher graph metrics than those without. These findings suggest the relative robustness of network segregation and small-worldness properties to denoising strategy. However, caution should be taken when interpreting path length findings across studies, as subtle variations in regression approach may account for discrepancies. Continued efforts should be taken towards harmonizing preprocessing pipelines across studies to aid replication efforts and build consensus towards understanding the mechanisms underlying pathological aging.

## Introduction

Alzheimer’s disease (AD) is an age-related neurodegenerative disorder characterized by the accumulation of two neuropathological hallmarks including amyloid-beta plaques (A*β*) and neurofibrillary tangles and progressive cognitive decline. AD neuropathology, particularly A*β*, develops years to decades prior to the onset of clinical symptoms (Jack Jr et al., 2018; Sperling et al., 2011), and as a consequence, alterations occur to structural and functional neural systems that underlie pathological cognitive decline. Though much knowledge has been garnered regarding the mechanisms underlying AD, the etiology and biological progression of AD remains elusive. As AD is projected to more than double by 2050 (Hebert, Weuve, Scherr, & Evans, 2013), a better understanding of the mechanisms underlying AD are needed to develop more targeted treatments and to identify biological markers to aid early intervention, when treatments may be most effective.

AD is proposed to be a disconnection syndrome (Delbeuck, Van der Linden, & Collette, 2003), where the precise spatiotemporal coherence of brain regions within and between neural networks is disrupted as a downstream consequence of neuropathology, which underpins cognition decline. One technique to better characterize network disruptions across the AD continuum is graph theoretical analysis of resting state fMRI. Graph theory models the brain and its functional connections as a network and examines the topology that underlies higher-order information processing and integration. A graph is a mathematical representation of a brain network and is constructed as a series of nodes and edges (Rubinov & Sporns, 2010). Nodes are comprised of a set of brain regions while edges represent the functional connection, or correlation, between the timeseries of each node. From these graphs, two global properties of the network can be explored including segregation and integration. Network segregation represents the ability for specialized processing to occur within clusters of highly connected regions of the network (Rubinov & Sporns, 2010). In functional networks, segregation properties describe the number of submodules present within the network exhibiting high intramodule connectivity. Network integration captures the ability of the network to rapidly transfer and combine information across specialized modules and distributed nodes (Rubinov & Sporns, 2010). Importantly, functional networks are shown to exhibit an optimal balance between segregation and integration, such that the network is organized to support subnetwork organization to perform specialized processing but also exhibits strategically placed links to facilitate efficient information integration across submodules. This network organizational efficiency can be quantified through small-worldness (Humphries & Gurney, 2008).

Graph theoretical analysis is frequently applied to study network alterations across the AD clinical continuum (Dai & He, 2014). This literature has revealed that pathological aging is associated with disruptions to both global network properties (segregation, integration) (Li et al., 2020; Brier et al., 2012; Si, Liu, Wang, Wang, & Zhao, 2019; Wang et al., 2013; Supekar, Menon, Rubin, Musen, & Greicius, 2008; Khazaee, Ebrahimzadeh, & Babajani-Feremi, 2016) and local nodal attributes (Subramanian et al., 2020; Behfar et al., 2020; Luo et al., 2021), features which can be used with high fidelity to distinguish cognitively normal (CN), MCI, and AD groups (J. Liu, Tan, Lan, & Wang, 2020; de Vos et al., 2018; Hojjati, Ebrahimzadeh, & Babajani-Feremi, 2019; Hojjati et al., 2017; Khazaee et al., 2016; Xu et al., 2020). A significant issue in this literature, however, is that there is little consensus in the reported direction of diagnostic group differences which impedes efforts to draw conclusions about the mechanisms underlying network dysfunction and their functional consequences in AD. For example, studies investigating clustering coefficient, a measure of network segregation, have reported higher clustering coefficient in MCI or AD than CN (Maulaz, de Almeida Mantovani, & da Silva, 2020; Zhao et al., 2012; Z. Liu et al., 2012; Dai et al., 2019), lower clustering coefficient in MCI or AD than CN (Xiang, Guo, Cao, Liang, & Chen, 2013; Si et al., 2019; Li et al., 2020; Brier et al., 2012; Dai & He, 2014; Xue et al., 2020, 2020), or no diagnostic differences (L. Zhang et al., 2020; Sanz-Arigita et al., 2010; Y. Liu et al., 2014; Khazaee et al., 2016). Similar discrepancies are reported with characteristic path length (Xiang et al., 2013; Zhao et al., 2012; Dai et al., 2019; Si et al., 2019; L. Zhang et al., 2020; Li et al., 2020; Brier et al., 2012; Supekar et al., 2008; Khazaee et al., 2016), a measure of network integration, and small-worldness (L. Zhang et al., 2020; Li et al., 2020; Dai et al., 2019; Supekar et al., 2008; Khazaee et al., 2016; Xue et al., 2020; Brier et al., 2012).

The lack of consensus may be due to a number of methodological factors including diagnostic criteria of MCI or AD patients included in the study, presence/load of AD neuropathology (A*β*) in each diagnostic group, functional data preprocessing pipeline, and node or edge definition (Dai & He, 2014). Here, we focus on the potential impact of preprocessing decisions, specifically denoising strategy, on AD diagnostic differences in graph metrics. An emerging literature provides evidence that the steps included and order of preprocessing pipelines affects the reliability of functional connectivity estimations and ultimately graph metrics (Alakörkkö, Saarimäki, Glerean, Saramäki, & Korhonen, 2017; Aurich, Alves Filho, Marques da Silva, & Franco, 2015; Borchardt et al., 2016; Liang et al., 2012; Ran et al., 2020; Vỳtvarová et al., 2017; Yan, Craddock, He, & Milham, 2013). For instance, Liang and colleagues (2012) found that preprocessing pipelines excluding global signal regression and including a wide bandpass frequency range (0.02-0.07) produced the most reliable topological estimates. Motion is particularly problematic in the context of fMRI, as sub-millimeter movements significantly distort functional connections between local and distant nodes (Power, Barnes, Snyder, Schlaggar, & Petersen, 2012; Van Dijk, Sabuncu, & Buckner, 2012). As such, the practice of censoring, or discarding high-motion volumes, has shown to be effective in addressing transient effects of motion in the BOLD signal (Power et al., 2014; Ciric et al., 2017) and lowers the correlation between motion and graph parameters, except when censoring is combined with global signal regression (Aurich et al., 2015). However, most previous work characterizing the influence of preprocessing steps on graph metrics have been employed in young, healthy cohorts (Alakörkkö et al., 2017; Aurich et al., 2015; Liang et al., 2012), and the impact of such decisions on diagnostic group differences have yet to be explored in the context of AD.

The purpose of the current study was to extend previous work by examining if diagnostic differences in graph metrics were dependent on denoising approach. Here, we focused on the combination of three preprocessing steps that are utilized to reduce the impact of physiological (i.e., bandpass filtering) and motion artefacts (i.e., temporal expansions of realignment parameters, censoring) on the BOLD signal. CN, MCI, and AD participants with resting-state fMRI and AV45 PET data were utilized from the Alzheimer’s Disease Neuroimaging Initiative (ADNI) database. Using AFNI, 16 preprocessing pipelines (Figure 1) were implemented to compare the impact of the combination of bandpass filtering (0.01-0.10 Hz), 12 motion parameters, censoring high-motion volumes, and censoring outlier volumes based on BOLD signal on diagnostic differences in graph metrics. Specifically, we calculated clustering coefficient, local efficiency, and modularity to quantify network segregation, largest connected component, characteristic path length and global efficiency to quantify network integration and small-worldness. We expected that metrics which have exhibited the most reported discrepancies in the AD literature (e.g., clustering coefficient, path length, and small-worldness) to show the highest dependency on denoising strategy.

**Figure 1.**
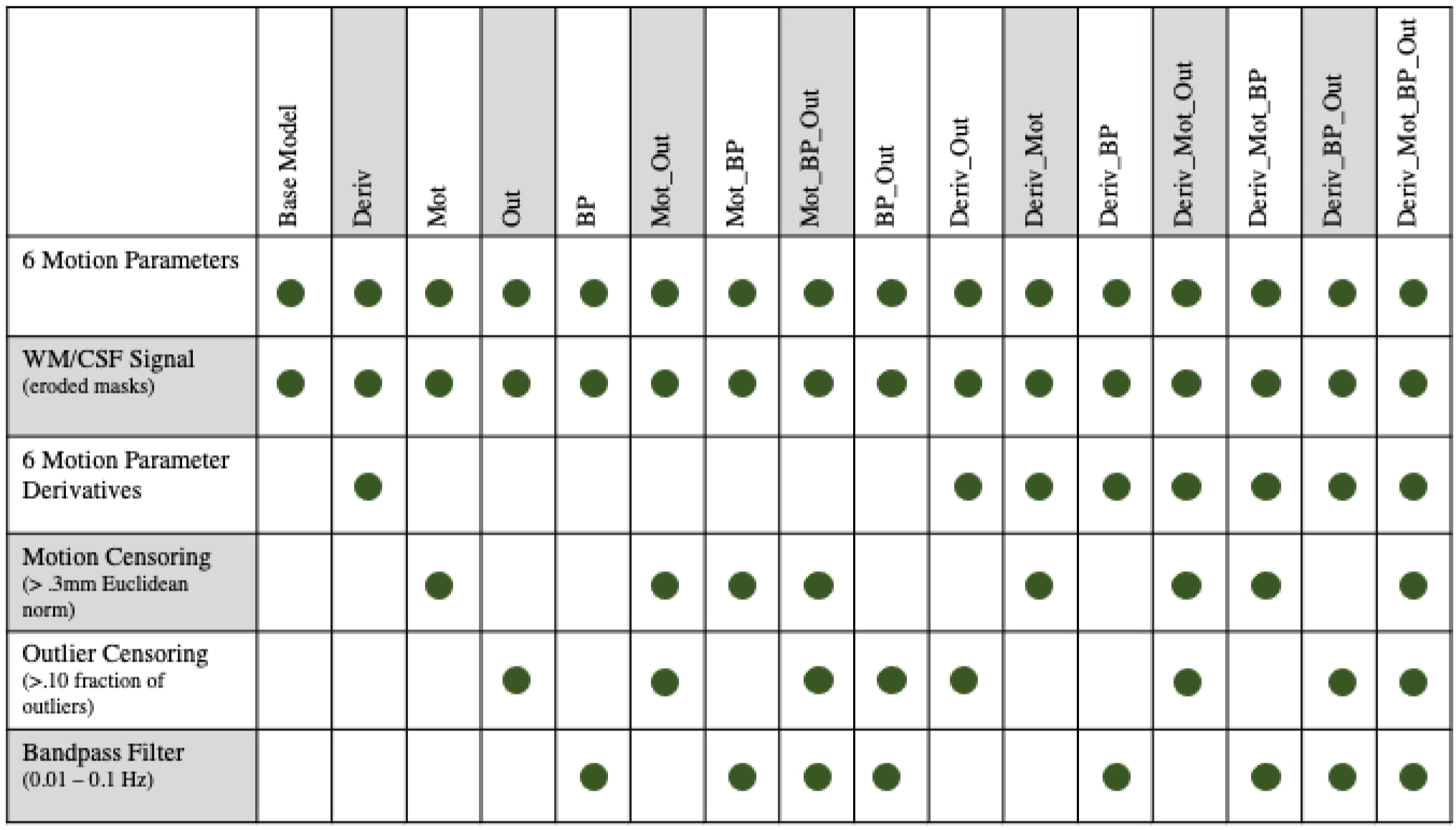
Sixteen nuisance regression models used for denoising. The denoising model is indicated at the top row of the figure. Green dots demonstrate which parameters were included in each model.

## Methods

### Participants

Data used in this study were obtained from the ADNI database (http://adni.loni.usc.edu). ADNI was launched in 2003 as a public-private partnership, led by Principal Investigator Michael W. Weiner, MD. The primary goal of ADNI has been to test whether serial magnetic resonance imaging (MRI), positron emission tomography (PET), other biological markers, and clinical and neuropsychological assessment can be combined to measure the progression of mild cognitive impairment (MCI) and early AD. For up-to-date information, see www.adni-info.org.

Participants were selected based on the availability of both resting-state fMRI and PET AV45 data. All PET scans occurred within one year of the rs-fMRI visit. The final sample included 60 cognitively normal older adults (CN), 60 MCI patients, and 40 AD patients matched for age and sex distribution (Table 1). PET data were utilized to determine amyloid-status for each individual with the criterion of 1.11 SUVR normalized to the cerebellum, as recommended by ADNI for cross-sectional investigations (Landau et al., 2012, 2013). Individuals with values above the criterion were deemed amyloid-positive and individuals below the threshold were deemed amyloid-negative (Jones et al., 2016). All MCI and AD participants were amyloid-positive while the amyloid-status for the CN group was mixed (Table 1). A list of participants and scans used in the current analysis can be found in Supplementary Table 1. The inclusion criteria for each diagnostic group were determined by ADNI. AD participants had a subjective memory concern, abnormal Logical Memory II subscale performance (≤8 for 16 or more years of education; ≤4 for 8-15 years of education; ≤2 for 0-7 years of education), Mini-Mental State Exam (MMSE) performance between 20-26, Clinical Dementia Rating (CDR) performance of 0.5 or 1.0, and meet the National Institute of Neurological and Communicative Disorders and Stroke and the AD and Related Disorders Association (NINCDS/ADRDA) criteria for probable AD. MCI participants had a subjective memory concern, MMSE performance of 24-30, CDR performance of 0.5, preserved functioning, and did not meet criteria for diagnosis of AD. CN participants were non-demented, non-MCI, free of memory complaints, had MMSE performance between 24-30, and CDR performance of 0.

**Table 1.** Sample Demographic Characteristics.

|  | CN | MCI | AD |
| --- | --- | --- | --- |
| N | 60 | 60 | 40 |
| Age (range) | 61.10-91.50 | 56.70-93.20 | 55.30-95.90 |
| Age | 75.29 (7.66) | 74.51 (7.85) | 76.87 (9.62) |
| Sex (% Female) | 53.33 | 48.33 | 50.00 |
| Education<sup>a</sup> | 16.88 (2.37) | 16.23 (2.56) | 15.20 (2.49) |
| APOE4 (% carriers)<sup>b</sup> | 28.33 | 58.33 | 72.50 |
| Amyloid-Positivity (%) | 43.33 | 100.00 | 100.00 |
| MMSE<sup>c</sup> | 29.12 (1.11) | 27.51 (2.12) | 21.85 (3.22) |
| Frame-to-Frame Motion | 0.09 (0.03) | 0.10 (0.04) | 0.11 (0.03) |
| Motion Censor Fraction | 0.05 (0.06) | 0.05 (0.05) | 0.07 (0.06) |
| Outlier Censor Fraction<sup>d</sup> | 0.01 (0.01) | 0.01 (0.02) | 0.02 (0.03) |
| Combined Censor Fraction | 0.05 (0.06) | 0.06 (0.05) | 0.08 (0.06) |

Note. Values presented as $M \pm SD$ unless otherwise specified.
<sup>a</sup>Significant main effect of diagnosis on education, $F(2,157) = 5.55$ , $p = .005$ . CN had more years of education than AD, $p = .003$ . There was a trend that MCI had more years of education than AD, $p = .072$ .
<sup>b</sup>Significant main effect of diagnosis on *APOE4* carrier distribution, $\chi^2(2) = 20.34$ , $p < .001$ . AD had a higher proportion of *APOE4* carriers than MCI and CN.
<sup>c</sup>Significant main effect of diagnosis on MMSE performance, $F(2,154) = 138.00$ , $p < .001$ . CN performed better on the MMSE than MCI and AD, $p$ 's $\leq .001$ . MCI performed better on the MMSE than AD, $p < .001$ .
<sup>d</sup>Significant main effect of diagnosis on outlier censor fraction, $F(2,157) = 4.35$ , $p = .014$ . There was a trend that AD had more volumes censored due to outliers than CN and MCI, $p$ 's $\leq .072$ .

### MRI acquisition and preprocessing

Imaging data from ADNI phases 2 and 3 (basic sequence) were used in the current analysis. The MR acquisition protocols employed across phases were common to all sites and adapted for different scanners. Functional images sensitized to BOLD contrast for rs-fMRI were collected on 3T scanners (Table 2). Participants were instructed to keep their eyes open during the rs-fMRI sequence.

**Table 2.**
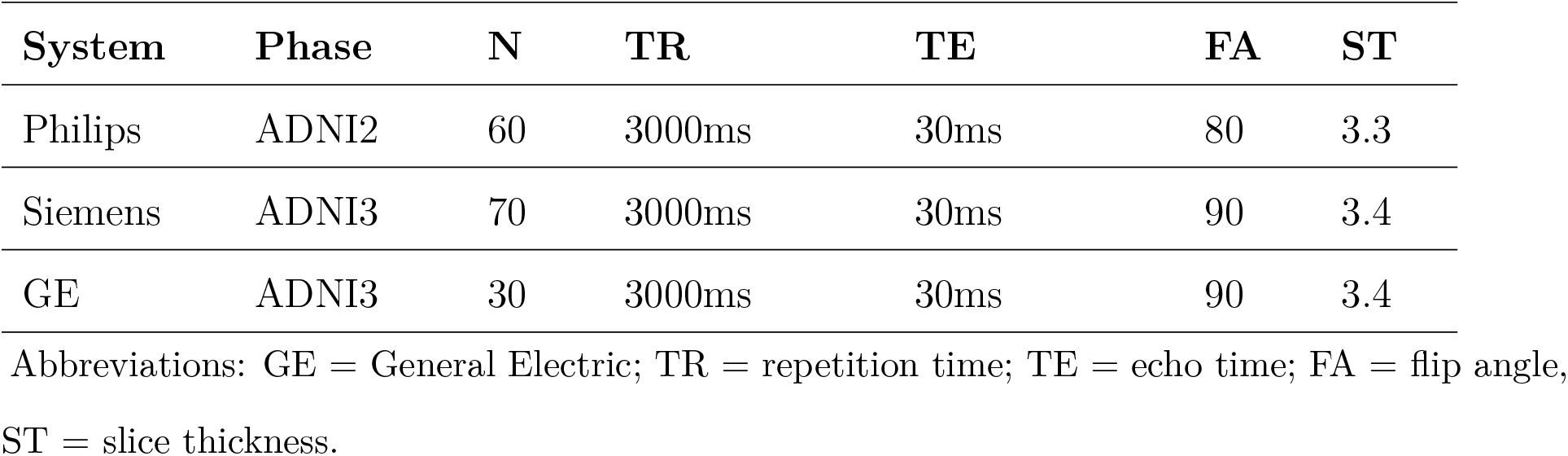
Functional rs-fMRI Acquisition Parameters by MRI Manufacturer.

Preprocessing was implemented using Freesurfer (v.7.1.1) (Fischl, 2012) and AFNI (v.21.2.04) (Cox, 1996). The centers of the anatomical and functional images were aligned to the center of a standard template prior to preprocessing (MNI152_2009_template_SSW; *Align_Centers*). Anatomical images were processed through AFNI’s *SSWarper* pipeline to skull strip the anatomy and calculate the warp to MNI space and Freesurfer’s *recon-all* pipeline to generate subject-specific WM and CSF masks.

AFNI’s *afni_proc*.*py* tool was used for functional preprocessing and included despike, tshift, align, tlrc, volreg, blur, mask, scale, and regress blocks. The first four volumes were removed to eliminate pre-steady state effects. Data were despiked (*3dDespike*) and the fraction of outliers within the brain mask were calculated for each volume (*3DToutcount*). The volume with the least outliers was used as the reference volume throughout processing. Datasets were slice-time corrected (*3dTshift*) according to two slice-time acquisitions patterns in the sample: altplus (N = 90) and alt+z2 (N = 70). Transforms for volume registration (*3dvolreg*), anatomy to functional (*align_epi_anat*.*py*), affine anatomy to MNI, and non-linear anatomy to MNI were concatenated and applied in a single step to minimize data interpolations. Data were smoothed with a FWHM kernel of 4mm (*3dmerge*). Voxel timeseries were scaled to a mean of 100 (*3dTstat/3dcalc*). A censor file was created per subject indicating volumes to be scrubbed where more than 10% of voxels were flagged as outliers and/or where frame-to-frame (Euclidean norm; enorm) motion was greater than 0.3mm (Grajski, Bressler, Initiative, et al., 2019; Weis et al., 2021)). All participants had less than 21% of volumes censored (Grajski et al., 2019). Diagnostic groups did not differ in overall levels of frame-to-frame motion, *F* (2, 157) = 2.23, *p* = .111 (Table 1).

### Nuisance regression approach

Sixteen preprocessing pipelines (Figure 1) were employed using each unique combination of nuisance regressors including (a) first temporal derivatives of the six realignment parameters (RP) (b) censoring due to motion (Euclidean norm (enorm) *>* 0.3mm) (c) censoring due to outliers (outlier fraction in volume *>* 0.10) and (d) bandpass filtering (0.01 - 0.10 Hz).

A. Base Model - Included six RP, local WM regressor, CSF signal, and linear and quadratic trends (polort=4) served as the base model for comparison in the current study.
B. Deriv - Base model + the first temporal derivatives of the six RP
C. Mot - Base model + motion censoring (volumes with enorm *>* 0.3mm)
D. Out - Base model + outlier censoring (volumes with more than 0.10 fraction of voxels identified as outliers)
E. BP - Base model + bandpass filtering (0.01 - 0.10 Hz)
F. BPOut - Base model + outlier censoring and bandpass filtering
G. MotOut - Base model + motion and outlier censoring
H. MotBP - Base model + motion censoring and bandpass filtering
I. MotBPOut - Base model + motion and outlier censoring and bandpass filtering
J. DerivMot - Base model + first temporal derivatives of six RP and motion censoring
K. DerivOut - Base model + first temporal derivatives of six RP and outlier censoring
L. DerivBP - Base model + first temporal derivatives of six RP and bandpass filtering
M. DerivMotOut - Base model + first temporal derivatives of six RP and motion and outlier censoring
N. DerivMotBP - Base model + first temporal derivatives of six RP, motion censoring, and bandpass filtering
O. DerivBPOut - Base model + first temporal derivatives of six RP, bandpass filtering, and outlier censoring
P. DerivMotBPOut - Base model + first temporal derivatives of six RP, motion censoring, outlier censoring, and bandpass filtering

### Nuisance Regressors

The base model comprised nuisance regressors commonly applied across age-related resting state investigations (Li et al., 2020; Si et al., 2019; Xiang et al., 2013). Six RP were derived from volume registration. Tissue-based regressors included ventricular CSF and localized WM signal. AFNI employs the ANATICOR approach to provide a localized WM regressor (Jo et al., 2013). This approach isolates spatially localized artefacts in WM, due to hardware or motion, that alter the fMRI time series (Caballero-Gaudes & Reynolds, 2017; Jo et al., 2013). The base model also included linear and quadratic detrending. The base model was employed in every iteration of pipelines tested.

The four regressors combined and iterated across pipelines included RP derivatives, bandpass filtering, motion censoring, and outlier censoring. The first temporal derivatives of the six RP were employed in nuisance regression to further reduce motion-related artefacts in the signal. The first temporal derivatives of the motion RP was employed in eight pipelines including Deriv, DerivOut, DerivMot, DerivBP, DerivMotOut, DerivMotBP, DerivBPOut, DerivMotBPOut.

Bandpass filtering was utilized to remove physiological (e.g., cardiac, respiratory) artefacts. A bandpass filter of 0.01 - 0.10 Hz was used in eight pipelines including BP, BPOut MotBP, MotBPOut, DerivBP, DerivMotBP, DerivBPOut, DerivMotBPOut.

Censoring, or scrubbing, was employed to further address residuals left in the data due to artefacts, such as motion or hardware, by completely removing the TR from analysis (Power et al., 2012). AFNI uses two approaches for censoring based on percentage of outliers and motion. Outlier voxels for each volume were calculated using AFNI’s *3doutcount* tool. Volumes to be censored due to outliers were designated as volumes where more than 10% of voxels are flagged as outliers. Outlier censoring was included in eight pipelines including Out, BPOut, MotOut, MotBPOut, DerivOut, DerivMotOut, DerivBPOut, and DerivMotBPOut. To detect volumes to be scrubbed due to motion, enorm was calculated for each volume, which estimated frame-to-frame motion. Volumes that exceeded enorm of 0.3mm were scrubbed from the time series (Grajski et al., 2019; Weis et al., 2021). Motion censoring (Mot) was included in eight pipelines: Mot, MotOut, MotBP, MotBPOut, DerivMot, DerivMotOut, DerivMotBP, and DerivMotBPOut. While the volumes flagged for censoring due to outliers largely overlap with volumes flagged for motion, outlier censoring may also capture non-motion-related residual artefacts, such as hardware.

### Graph Construction

The brain was parcellated using 79 functional ROIs (nodes; Figure 2) defined across 14 resting state networks affected in AD (Shirer, Ryali, Rykhlevskaia, Menon, & Greicius, 2012). Regions encompassed by each node are described in Supplemental Table 2. This atlas provided comprehensive coverage of the cortical and subcortical regions. The average time series from each node was Pearson correlated and Fischer z-transformed to yield a 79×79 functional connectivity matrix per individual. Currently, there is no gold-standard in the field on how to handle negative weights in the correlation matrix (Hallquist & Hillary, 2018; Rubinov & Sporns, 2010). Two common approaches described in the literature include absolutizing the correlation coefficients or retaining only positive correlations by zeroing out negative correlations in the matrix (Hallquist & Hillary, 2018; Ran et al., 2020). Past work suggests that negative correlations contain biologically relevant information that can aid group classification (Kazeminejad & Sotero, 2020) and retaining information from negative correlations through absolute values improves the reproducibility of graph metrics (Ran et al., 2020). Thus, information from negative correlations were retained in the current study by taking the absolute value of the correlation matrix (de Vos et al., 2018; Schwarz & McGonigle, 2011). The absolutized connectivity matrix was transformed into a binary adjacency matrix (graph) for calculation of graph properties by setting all self-connections (diagonal) to 0, thresholding, and binarizing such that all subthreshold values were set to 0 and all suprathreshold values set to 1. A proportional thresholding procedure was utilized to make certain that the same density of connections (i.e., edges in the graph) were retained across individuals. To ensure that group differences in graph properties were not dependent on graph density, the matrices were proportionally thresholded across a wide range of densities, retaining the top 2.5% to 25% of edges at steps of 2.5 (Langella, Sadiq, Mucha, Giovanello, & Dayan, 2021), and graph metrics were calculated at each density. For ease of interpretation, graph properties were averaged across all densities.

**Figure 2.**
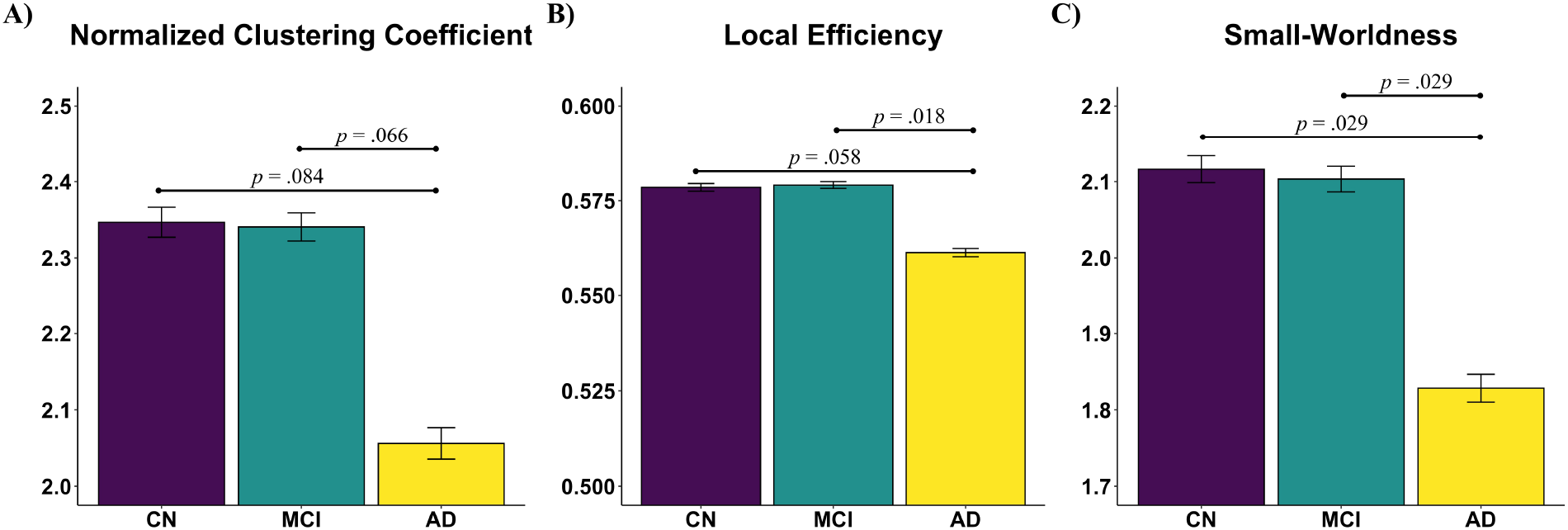
Diagnostic differences in graph metrics. (A) Clustering coefficient, (B) local efficiency, (C) and small-worldness significantly differed by diagnostic group, such that CN and MCI had greater network segration and small-worldness than AD.

### Graph Metrics

Graph metrics were calculated using the Brain Connectivity Toolbox (Rubinov & Sporns, 2010). Six graph measures were used to characterize network segregation and network integration. Segregation properties were quantified through clustering coefficient, modularity, and local efficiency metrics. Normalized clustering coefficient (CC_real_/CC_rand_) measures the fraction of neighbors of a node that are neighbors of each other (Watts & Strogatz, 1998). CC_real_ was quantified from the brain network. CC_real_ was normalized to CC_rand_ calculated from 30 random networks with the same density of the real network where each edge was rewired approximately 10 times (C. Zhang et al., 2016). Modularity quantifies the number of submodules with maximal intramodule and minimal intermodule connections that the overall network can be separated into (Newman, 2004). Local efficiency represents the efficiency of information transfer within a neighborhood of nodes (Latora & Marchiori, 2001). Network integration in the current study was captured through largest connected component, characteristic path length, and global efficiency metrics. Largest connected component is a metric that serves as a prerequisite of integration and quantifies node connectivity within the graph. Graphs which have larger connected components (i.e., higher nodal connectivity) are more likely to have efficient integration properties. Normalized path length (PL_real_/PL_rand_) represents the shortest average path length between any two nodes in the graph (Watts & Strogatz, 1998). Global efficiency was calculated as the inverse of the shortest path length (Latora & Marchiori, 2001). Small-worldness was calculated as the ratio of normalized clustering coefficient to normalized path length. Values greater than 1 represent small-world architecture.

### Statistical Analysis

Statistical analyses were conducted using R (4.1.0). Diagnostic differences in demographic variables were calculated using the *ezANOVA* function from the *ez* package. To examine the effect of diagnostic status and nuisance regression approach on graph properties, a series of 3 × 16 mixed ANCOVA models were calculated using the *aov_car* function of the *afex* package. Age, sex, education, and *APOE4* genotype were included in each model as covariates. In circumstances where sphericity assumptions were violated, Greenhouse-Geiser corrections were applied. Posthoc comparisons were conducted using *emmeans*. Correction for multiple comparisons were implemented using the Tukey method.

## Results

### Demographics

Detailed demographics are presented in Table 1. Diagnostic groups did not differ in terms of age, *F* (2, 157) = 0.99, *p* = .374, or sex distribution, *χ*^2^(2) = 0.31, *p* = 0.86. There was a significant main effect of diagnosis on years of education, *F* (2, 157) = 5.56, *p* = .005. CN had more years of education than AD, *p* = .003. There was a trend that MCI had more years of education than AD, *p* = .072. There was a main effect of diagnosis on MMSE performance, *F* (2, 154) = 138.00, *p <* .001. CN and MCI performed better on the MMSE than AD, *p*’s *<* .001, and CN performed better on the MMSE than MCI, *p <* .001. There was a significant relationship between diagnostic group and *APOE4* genotype, *χ*^2^(2) = 20.34, *p <* .001.

### Network Segregation

#### Clustering Coefficient

There was a trending main effect of diagnosis on clustering coefficient, *F* (2, 152) = 2.99, *p* = .053 (Figure 2A). CN and MCI had higher clustering coefficient than AD, *p*’s ≤ .084. There was a significant main effect of nuisance regression approach on clustering coefficient, *F* (3.19, 485.20) = 7.80, *p <* .001 (Figure 3A). Pipelines BP, BPOut, and MotBP produced higher clustering coefficient than Out, *p ≤* .047. MotBPOut produced higher clustering coefficient than the base model, Out, MotOut, and DerivMot, *p ≤* .041. DerivBP, DerivMotBP, DerivBPOut, and DerivMotBPOut pipelines produced higher clustering coefficient than the base model, Deriv, Mot, Out, MotOut, DerivMot, DerivOut, and DerivMotOut, *p ≤* .039. The interaction between diagnosis and nuisance approach was not significant, *F* (6.38, 485.2) = 0.55, *p* = .781.

**Figure 3.**
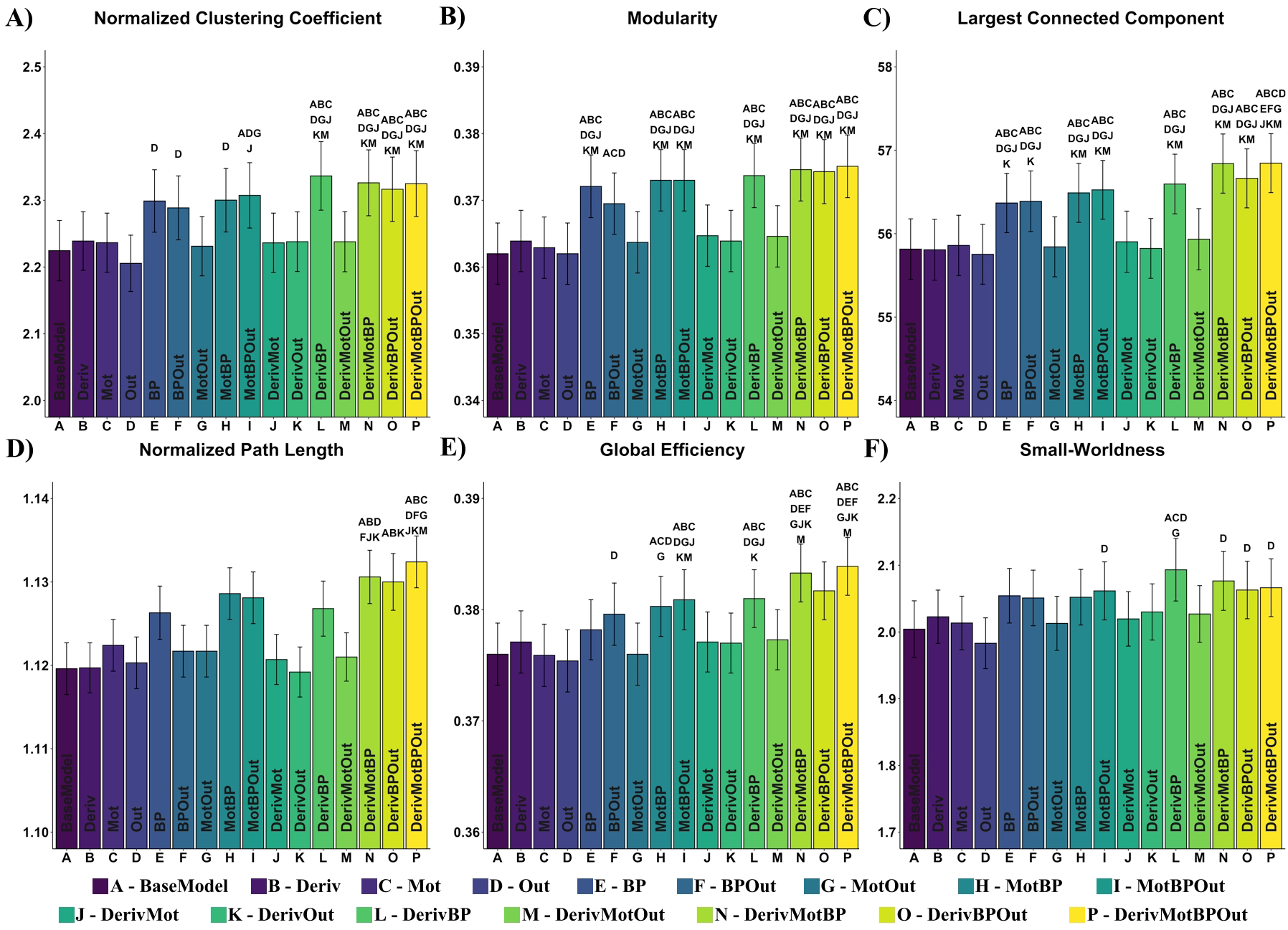
Significant main effects of nuisance regression approach on graph metrics. Nuisance models are indicated by letter with a full description of the model in each bar. Significant posthoc comparisons (after Tukey correction; p < .05) are denoted above each bar describing which models (designated by letters A-P) significantly differed in the magnitude of (A) clustering coefficient (B) modularity (C) largest connected component (D) path length (E) global efficiency, and (F) small-worldness. Generally, models including bandpass filtering produced higher graph metrics than those without.

**Figure 4.**
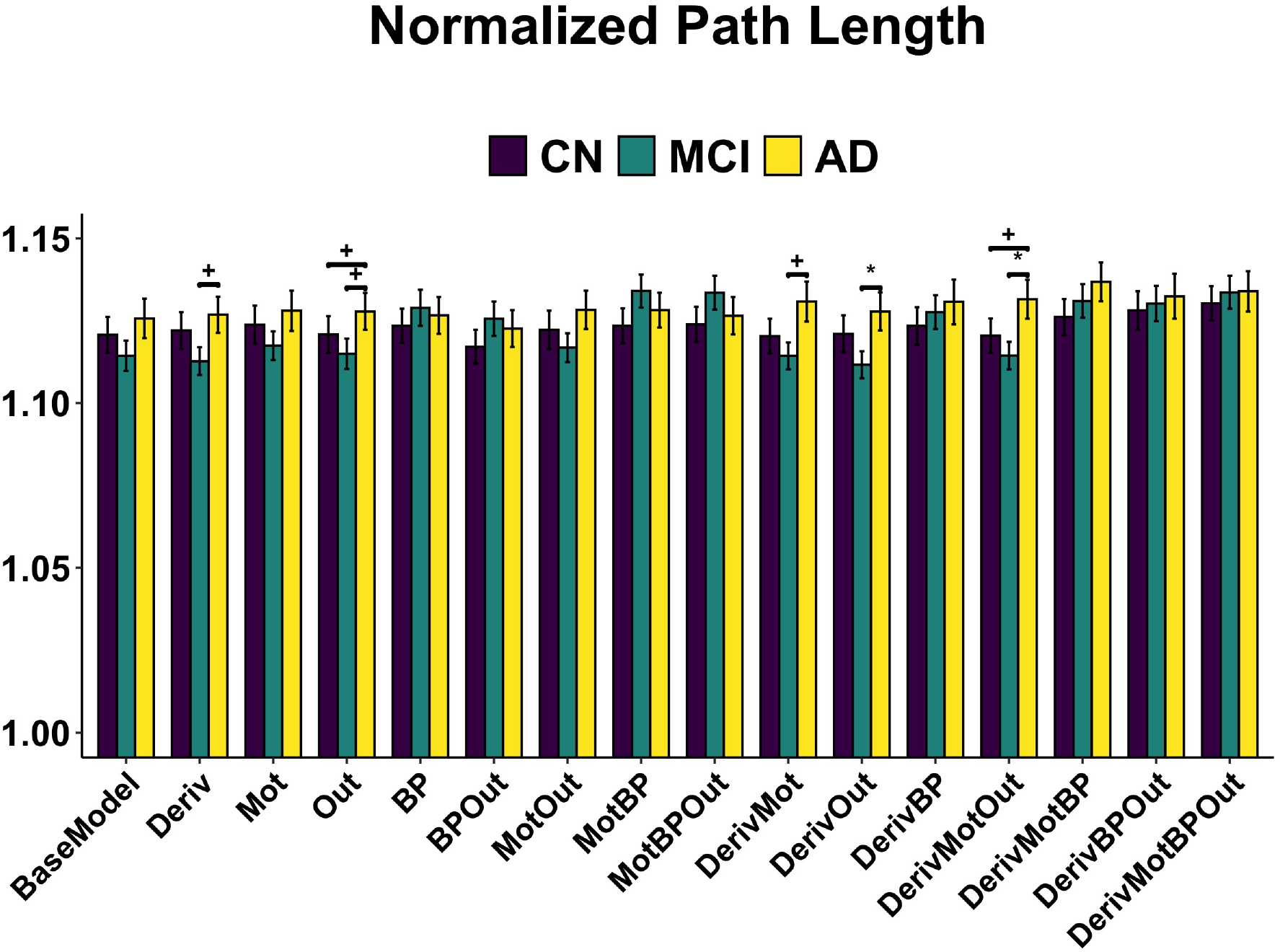
Significant interaction between diagnostic group and nuisance approach on path length. Diagnostic differences in path length, such that CN and MCI had lower, more efficent path lengths, than AD were only evident in Deriv, Out, DerivMot, DerivOut, and DerivMotOut pipelines. ^*^p’s ≤ .05, ^+^p’s ≤ .1

#### Modularity

The main effect of diagnosis on modularity was not significant, *F* (2, 152) = 2.00, *p* = .139. There was a significant main effect of nuisance approach on modularity, *F* (2.56, 389.36) = 14.45, *p <* .001 (Figure 3B). The BPOut pipeline produced higher modularity than the base model, Mot, and Out pipelines, *p*’s ≤ .020. The BP, MotBP, DerivBP, DerivMotBP, DerivBPOut, MotBPOut, and DerivMotBPOut pipelines output higher modularity than the base model, Deriv, Mot, Out, MotOut, DerivMot, DerivOut, and DerivMotOut pipelines, *p*’s ≤ .009. The interaction effect of diagnosis and nuisance approach on modularity was not significant, *F* (5.12, 389.36) = 0.32, *p* = .903.

#### Local Efficiency

There was a main effect of diagnosis on local efficiency, *F* (2, 152) = 4.07, *p* = .019 (Figure 2B). MCI had higher local efficiency than AD, *p* = .018. There was a trend that CN had higher local efficiency than AD, *p* = .058. The main effect of nuisance regression approach, *F* (2.81, 427.18) = 0.99, *p* = .392, and interaction effect, *F* (5.62, 427.18) = 0.75, *p* = .601, were not significant.

### Network Integration

#### Largest Connected Component

The main effect of diagnosis on largest connected component was not significant, *F* (2, 152) = 1.64, *p* = .197. There was a significant main effect of nuisance regression approach on largest connected component, *F* (2.73, 414.80) = 17.90, *p <* .001 (Figure 3C). DerivMotBPOut produced larger connected components than BP and BPOut, *p ≤* .045. The BP and BPOut pipelines showed larger connected components than the base model, Deriv, Mot, Out, MotOut, DerivMot, and DerivOut pipelines, *p*’s ≤ .030. The DerivBP, MotBP, MotBPOut, DerivMotBP, DerivBPOut, and DerivMotBPOut models produced larger connected components than the base model, Deriv, Mot, Out, MotOut, DerivMot, DerivOut, and DerivMotOut pipelines, *p*’s ≤ .002. The interaction effect of diagnosis and nuisance approach on largest connected component was not significant, *F* (5.46, 414.80) = 0.71, *p* = .626.

#### Characteristic Path Length

The main effect of diagnosis on path length was not significant, *F* (2, 152) = 1.56, *p* = .214. There was a significant main effect of nuisance pipeline on path length, *F* (3.76, 570.93) = 4.93, *p ≤* .001 (Figure 3D), and a significant interaction between diagnosis and nuisance approach on path length, *F* (7.51, 570.93) = 2.39, *p* = .018. MCI exhibited lower, more efficient path lengths than AD on DerivOut and DerivMotOut pipelines, *p*’s ≤ .043. There was a non-significant trend that CN had lower path lengths than AD on Out and DerivMotOut pipelines, *p* = .095. There was a non-significant trend that MCI had lower path lengths than AD on Deriv, Out, and DerivMot pipelines, *p*’s ≤ .079. Within CN, path length was significantly higher for DerivMotBPOut than BPOut, *p* = .012. Within MCI, MotBP and MotBPOut exhibited higher path length than the base model, Deriv, Out, DerivMot, DerivOut, and DerivMotOut pipelines, *p*’s ≤ .035. MCI levels further varied across pipelines with path length being higher in DerivMotBPOut than the base model, Deriv, DerivMot, DerivOut, DerivMotOut, *p*’s ≤ .048. Within MCI, path length was higher for DerivMotBP and DerivBPOut than Deriv and DerivOut pipelines, *p*’s ≤ .033. Within AD, the levels of path length did not significantly differ across denoising pipelines, *p*’s *≥* .463.

#### Global Efficiency

The main effect of diagnosis on global efficiency was not significant, *F* (2, 152) = 0.85, *p* = .429. There was a significant main effect of regression approach on global efficiency, *F* (2.42, 368.15) = 12.93, *p <* .001 (Figure 3E). BPOut produced higher global efficiency than Out, *p* = .047. The DerivMotBP and DerivMotBPOut pipelines produced higher global efficiency than BP and BPOut, *p*’s ≤ .018. MotBP showed higher global efficiency than the base model, Mot, Out, and MotOut, *p*’s ≤ .005. The DerivBP pipeline produced higher global efficiency than the base model, Deriv, Mot, Out, MotOut, DerivMot, and DerivOut pipelines, *p*’s ≤ .035. The MotBPOut, DerivMotBP, DerivBPOut, and DerivMotBPOut pipelines produced higher global efficiency than the base model, Deriv, Mot, Out, MotOut, DerivMot, DerivOut, and DerivMotOut pipelines, *p*’s ≤ .043. The interaction effect of diagnosis and nuisance approach on global efficiency was not significant, *F* (4.84, 368.15) = 0.90, *p* = .478.

### Small-Worldness

There was a significant main effect of diagnosis on small-worldness, *F* (2, 152) = 4.12, *p* = .018 (Figure 2C). AD had lower small-worldness than CN and MCI, *p*’s ≤ .029. There was a significant main effect of nuisance regression approach on small-worldness, *F* (3.72, 564.97) = 3.54, *p* = .009 (Figure 3F). MotBPOut, DerivMotBP, DerivBPOut, and DerivMotBPOut had higher small-worldness than Out, *p*’s ≤ .042. DerivBP had higher small-worldness than the base model, Mot, Out, and MotOut, *p*’s ≤ .037. The interaction of diagnosis and nuisance regression approach was not significant, *F* (7.43, 564.97) = 1.53, *p* = .151.

## Discussion

Graph theoretical analysis offers a unique lens to identify alterations in network topology that underlie pathological cognitive decline. While past work in the context of AD has revealed disrupted network integration, segregation, and small-worldness (Li et al., 2020; Brier et al., 2012; Supekar et al., 2008; Khazaee et al., 2016), the directionality of these differences across diagnostic groups varies by study, limiting the ability to draw concordant conclusions on the brain mechanisms underlying pathological aging. The analysis pipeline used to generate graph metrics is multifaceted, including steps such as fMRI preprocessing/denoising, the use or size of smoothing kernel, brain parcellation scheme, and definition of edges (e.g., correlation technique, thresholding criteria, weighting) (Farahani, Karwowski, & Lighthall, 2019; Hallquist & Hillary, 2018). As there is currently no gold-standard in the field for each of these decision-points, the collective pipeline can vary significantly from study to study (Hallquist & Hillary, 2018), likely contributing to discrepant reportings. The purpose of the current investigation was to determine if diagnostic differences in functional network topology were dependent on denoising approach. Specifically, comparisons of graph metrics generated across 16 pipelines that iterated combinations of denoising approach were made (e.g., bandpass filtering, temporal expansion of six realignment parameters, censoring due to motion, censoring due to outliers). Three main findings emerged: (1) diagnostic differences in characteristic path length, one of the primary measures used to characterize network integration, was dependent on denoising strategy; (2) diagnostic group differences in network segregation properties, as measured via clustering coefficient and local efficiency, and small-worldness were robust to denoising approach; (3) regardless of diagnostic group, denoising strategy influenced the magnitude of graph metrics, such that pipelines including bandpass filtering exhibited higher graph properties than pipelines excluding this denoising step.

Much work documenting the influence of denoising strategy on network topology has been conducted in young healthy adults (Aurich et al., 2015; Liang et al., 2012; Ciric et al., 2017; Vỳtvarová et al., 2017; Yan et al., 2013). A strength of the current investigation was that analyses were conducted in older adults at different stages of the AD continuum. In line with past work (Aurich et al., 2015), we observed that the inclusion of bandpass filtering into the denoising pipeline significantly influenced all graph metrics except local efficiency. Temporal filtering is typically employed in resting state preprocessing to remove physiological (i.e., cardiac, respiratory) and motion-related noise components from the BOLD signal (Caballero-Gaudes & Reynolds, 2017; Hallquist, Hwang, & Luna, 2013). Higher magnitude of graph metrics in pipelines including bandpass filtering observed in the current study may reflect reduced “non-neuronal” noise within the signal. However, future work is needed to discern the mathematical underpinnings of the observed bandpass filter-related effect.

In addition to the main effects of denoising approach on graph metrics, our results showed that the effect of denoising strategy on characteristic path length, in particular, was not ubiquitous across diagnostic groups, such that path length in the MCI and CN were most variable in pipelines including bandpass filtering. Despite the relative success of simultaneous bandpass filtering and nuisance regression to reduce noise, past work demonstrates that motion-related hyperconnectivity of short-range connections can persist (Hallquist et al., 2013). One possibility is that the bandpass filter-related alterations in graph metric magnitudes may reflect motion differences between groups. However, this is unlikely in the current study, as groups were matched for frame-to-frame motion (Table 1). It will be important in future studies to discern the variables driving diagnostic differences in bandpass-related variability in characteristic path length, given that path length is one of the most commonly used metrics to quantify network integration, and is used to calculate small-worldness, a network property affected in AD. If, for example, motion-related differences between groups are exacerbated by including bandpass filtering in pipelines, this is a critical methodological obstacle to consider when composing a processing pipeline and interpreting results.

Inconsistent with our hypothesis, we found that diagnostic differences in small-worldness and certain measures of network segregation, including clustering coefficient and local efficiency, were robust to denoising approach. This result likely suggests that the reported discrepancies in AD-related directionality in these metrics (Z. Liu et al., 2012; Brier et al., 2012; Li et al., 2020; L. Zhang et al., 2020; Sanz-Arigita et al., 2010; Khazaee et al., 2016; Si et al., 2019; Supekar et al., 2008) are not due to slight variations in denoising approach but rather reflect differences in decision-points downstream of denoising in the graph analysis pipeline. For example, it has been shown that there is a wide variety of atlases used to parcellate the functional brain across studies (Hallquist & Hillary, 2018) and that the network topology derived from different parcellation schemes significantly varies (Wang et al., 2009). Moreover, though most graph theory studies in the context of pathological aging utilize binarized graphs to estimate topology, there is significant variance across studies in the thresholding schemes utilized to define what constitutes a connection in a binarized network (e.g., proportional thresholding, absolute thresholding). Thus, more work is needed to investigate how differences in graph formation (node and edge definition) contribute to the discrepancies in AD-related alterations to network topology across the continuum.

Several limitations of the current study should be considered. First, though efforts were made to ensure that preprocessing steps, and especially denoising strategies, were not software-specific, certain features of the preprocessing pipeline were unique to AFNI, including despiking. It was ultimately decided to include despiking into the pipeline based on the evidence that despiking is effective at tempering the effects of motion artifacts (Jo et al., 2013), a particularly deleterious problem in resting state fMRI, and because the implementation of despiking has become accessible through programs like fMRIPrep (Esteban et al., 2019) in circumstances where users are not solely preprocessing functional data using one software (e.g., AFNI). A second limitation of the study is the focus on a small number of denoising approaches (e.g., first temporal derivatives of six realignment parameters, motion censoring, outlier censoring, and bandpass filtering). Past work has documented the influence of a multitude of denoising strategies on functional connectivity (Ciric et al., 2017; Power et al., 2014; Jo et al., 2013), such as noise removal via independent components analysis, and future work should examine the effect of these approaches on an array of graph metrics. The limitations of this work, however, should not underscore its strengths, including the investigation of the consequences of systematic methodological manipulations on network topology in a pathologically-characterized (.e.g, A*β* deposition) sample of older adults.

In summary, our results suggest that diagnostic differences in path length, but not clustering coefficient, local efficiency, or small-worldness, are susceptible to the implemented denoising strategy. It is critical that the field continues to practice transparent reporting of the methodologies implemented for processing and denoising of fMRI data to aid replication efforts and synthesis of findings across studies. Further, continued efforts should be taken towards identifying best practices, if possible, and harmonizing processing pipelines across studies to build consensus towards understanding the mechanisms underlying pathological aging.

## Acknowledgments

Data collection and sharing for this project was funded by the Alzheimer’s Disease Neuroimaging Initiative (ADNI) (National Institutes of Health Grant U01 AG024904) and DOD ADNI (Department of Defense award number W81XWH-12-2-0012). ADNI is funded by the National Institute on Aging, the National Institute of Biomedical Imaging and Bioengineering, and through generous contributions from the following: AbbVie, Alzheimer’s Association; Alzheimer’s Drug Discovery Foundation; Araclon Biotech; BioClinica, Inc.; Biogen; Bristol-Myers Squibb Company; CereSpir, Inc.; Cogstate; Eisai Inc.; Elan Pharmaceuticals, Inc.; Eli Lilly and Company; EuroImmun; F. Hoffmann-La Roche Ltd and its affiliated company Genentech, Inc.; Fujirebio; GE Healthcare; IXICO Ltd.; Janssen Alzheimer Immunotherapy Research & Development, LLC.; Johnson & Johnson Pharmaceutical Research & Development LLC.; Lumosity; Lundbeck; Merck & Co., Inc.; Meso Scale Diagnostics, LLC.; NeuroRx Research; Neurotrack Technologies; Novartis Pharmaceuticals Corporation; Pfizer Inc.; Piramal Imaging; Servier; Takeda Pharmaceutical Company; and Transition Therapeutics. The Canadian Institutes of Health Research is providing funds to support ADNI clinical sites in Canada. Private sector contributions are facilitated by the Foundation for the National Institutes of Health (www.fnih.org). The grantee organization is the Northern California Institute for Research and Education, and the study is coordinated by the Alzheimer’s Therapeutic Research Institute at the University of Southern California. ADNI data are disseminated by the Laboratory for Neuro Imaging at the University of Southern California. This reserach was additionally supported by NIH R01AG068990 (H. Oh), NIH R01AG069265 (W. Heindel/S. Buka), and NIH S10OD025181 (J. Sanes).

## References

Alakörkkö, T., Saarimäki, H., Glerean, E., Saramäki, J., & Korhonen, O. (2017). Effects of spatial smoothing on functional brain networks. European Journal of Neuroscience, 46 (9), 2471–2480.

Aurich, N. K., Alves Filho, J. O., Marques da Silva, A. M., & Franco, A. R. (2015). Evaluating the reliability of different preprocessing steps to estimate graph theoretical measures in resting state fmri data. Frontiers in neuroscience, 9, 48.

Behfar, Q., Behfar, S. K., Von Reutern, B., Richter, N., Dronse, J., Fassbender, R., … Onur, O. A. (2020). Graph theory analysis reveals resting-state compensatory mechanisms in healthy aging and prodromal alzheimer’s disease. Frontiers in aging neuroscience, 355.

Borchardt, V., Lord, A. R., Li, M., van der Meer, J., Heinze, H.-J., Bogerts, B., … Walter, M. (2016). Preprocessing strategy influences graph-based exploration of altered functional networks in major depression. Human brain mapping, 37 (4), 1422–1442.

Brier, M. R., Thomas, J. B., Snyder, A. Z., Benzinger, T. L., Zhang, D., Raichle, M. E., … Ances, B. M. (2012). Loss of intranetwork and internetwork resting state functional connections with alzheimer’s disease progression. Journal of Neuroscience, 32 (26), 8890–8899.

Caballero-Gaudes, C., & Reynolds, R. C. (2017). Methods for cleaning the bold fmri signal. Neuroimage, 154, 128–149.

Ciric, R., Wolf, D. H., Power, J. D., Roalf, D. R., Baum, G. L., Ruparel, K., … others (2017). Benchmarking of participant-level confound regression strategies for the control of motion artifact in studies of functional connectivity. Neuroimage, 154, 174–187.

Cox, R. W. (1996). Afni: software for analysis and visualization of functional magnetic resonance neuroimages. Computers and Biomedical research, 29 (3), 162–173.

Dai, Z., & He, Y. (2014). Disrupted structural and functional brain connectomes in mild cognitive impairment and alzheimer’s disease. Neuroscience Bulletin, 30 (2), 217–232.

Dai, Z., Lin, Q., Li, T., Wang, X., Yuan, H., Yu, X., … Wang, H. (2019). Disrupted structural and functional brain networks in alzheimer’s disease. Neurobiology of aging, 75, 71–82.

Delbeuck, X., Van der Linden, M., & Collette, F. (2003). Alzheimer’disease as a disconnection syndrome? Neuropsychology review, 13 (2), 79–92.

de Vos, F., Koini, M., Schouten, T. M., Seiler, S., van der Grond, J., Lechner, A., … Rombouts, S. A. (2018). A comprehensive analysis of resting state fmri measures to classify individual patients with alzheimer’s disease. Neuroimage, 167, 62–72.

Esteban, O., Markiewicz, C. J., Blair, R. W., Moodie, C. A., Isik, A. I., Erramuzpe, A., … others (2019). fmriprep: a robust preprocessing pipeline for functional mri. Nature methods, 16 (1), 111–116.

Farahani, F. V., Karwowski, W., & Lighthall, N. R. (2019). Application of graph theory for identifying connectivity patterns in human brain networks: a systematic review. frontiers in Neuroscience, 13, 585.

Fischl, B. (2012). Freesurfer. Neuroimage, 62 (2), 774–781.

Grajski, K. A., Bressler, S. L., Initiative, A. D. N., et al. (2019). Differential medial temporal lobe and default-mode network functional connectivity and morphometric changes in alzheimer’s disease. NeuroImage: Clinical, 23, 101860.

Hallquist, M. N., & Hillary, F. G. (2018). Graph theory approaches to functional network organization in brain disorders: A critique for a brave new small-world. Network Neuroscience, 3 (1), 1–26.

Hallquist, M. N., Hwang, K., & Luna, B. (2013). The nuisance of nuisance regression: spectral misspecification in a common approach to resting-state fmri preprocessing reintroduces noise and obscures functional connectivity. Neuroimage, 82, 208–225.

Hebert, L. E., Weuve, J., Scherr, P. A., & Evans, D. A. (2013). Alzheimer disease in the united states (2010–2050) estimated using the 2010 census. Neurology, 80 (19), 1778–1783.

Hojjati, S. H., Ebrahimzadeh, A., & Babajani-Feremi, A. (2019). Identification of the early stage of alzheimer’s disease using structural mri and resting-state fmri. Frontiers in neurology, 904.

Hojjati, S. H., Ebrahimzadeh, A., Khazaee, A., Babajani-Feremi, A., Initiative, A. D. N., et al. (2017). Predicting conversion from mci to ad using resting-state fmri, graph theoretical approach and svm. Journal of neuroscience methods, 282, 69–80.

Humphries, M. D., & Gurney, K. (2008). Network ‘small-world-ness’: a quantitative method for determining canonical network equivalence. PloS one, 3 (4), e0002051.

Jack Jr, C. R., Bennett, D. A., Blennow, K., Carrillo, M. C., Dunn, B., Haeberlein, S. B., … others (2018). Nia-aa research framework: toward a biological definition of alzheimer’s disease. Alzheimer’s & Dementia, 14 (4), 535–562.

Jo, H. J., Gotts, S. J., Reynolds, R. C., Bandettini, P. A., Martin, A., Cox, R. W., & Saad, Z. S. (2013). Effective preprocessing procedures virtually eliminate distance-dependent motion artifacts in resting state fmri. Journal of applied mathematics, 2013.

Jones, D. T., Knopman, D. S., Gunter, J. L., Graff-Radford, J., Vemuri, P., Boeve, B. F., … Jack Jr, C. R. (2016). Cascading network failure across the alzheimer’s disease spectrum. Brain, 139 (2), 547–562.

Kazeminejad, A., & Sotero, R. C. (2020). The importance of anti-correlations in graph theory based classification of autism spectrum disorder. Frontiers in neuroscience, 14, 676.

Khazaee, A., Ebrahimzadeh, A., & Babajani-Feremi, A. (2016). Application of advanced machine learning methods on resting-state fmri network for identification of mild cognitive impairment and alzheimer’s disease. Brain imaging and behavior, 10 (3), 799–817.

Landau, S. M., Lu, M., Joshi, A. D., Pontecorvo, M., Mintun, M. A., Trojanowski, J. Q., … Initiative, A. D. N. (2013). Comparing positron emission tomography imaging and cerebrospinal fluid measurements of β-amyloid. Annals of neurology, 74 (6), 826–836.

Landau, S. M., Mintun, M. A., Joshi, A. D., Koeppe, R. A., Petersen, R. C., Aisen, P. S., … Initiative, A. D. N. (2012). Amyloid deposition, hypometabolism, and longitudinal cognitive decline. Annals of neurology, 72 (4), 578–586.

Langella, S., Sadiq, M. U., Mucha, P. J., Giovanello, K. S., & Dayan, E. (2021). Lower functional hippocampal redundancy in mild cognitive impairment. Translational Psychiatry, 11 (1), 1–12.

Latora, V., & Marchiori, M. (2001). Efficient behavior of small-world networks. Physical review letters, 87 (19), 198701.

Li, W., Wen, W., Chen, X., Ni, B., Lin, X., Fan, W., … others (2020). Functional evolving patterns of cortical networks in progression of alzheimer’s disease: a graph-based resting-state fmri study. Neural Plasticity, 2020.

Liang, X., Wang, J., Yan, C., Shu, N., Xu, K., Gong, G., & He, Y. (2012). Effects of different correlation metrics and preprocessing factors on small-world brain functional networks: a resting-state functional mri study. PloS one, 7 (3), e32766.

Liu, J., Tan, G., Lan, W., & Wang, J. (2020). Identification of early mild cognitive impairment using multi-modal data and graph convolutional networks. BMC bioinformatics, 21 (6), 1–12.

Liu, Y., Yu, C., Zhang, X., Liu, J., Duan, Y., Alexander-Bloch, A. F., … Bullmore, E. (2014). Impaired long distance functional connectivity and weighted network architecture in alzheimer’s disease. Cerebral Cortex, 24 (6), 1422–1435.

Liu, Z., Zhang, Y., Yan, H., Bai, L., Dai, R., Wei, W., … others (2012). Altered topological patterns of brain networks in mild cognitive impairment and alzheimer’s disease: a resting-state fmri study. Psychiatry Research: Neuroimaging, 202 (2), 118–125.

Luo, Y., Sun, T., Ma, C., Zhang, X., Ji, Y., Fu, X., & Ni, H. (2021). Alterations of brain networks in alzheimer’s disease and mild cognitive impairment: A resting state fmri study based on a population-specific brain template. Neuroscience, 452, 192–207.

Maulaz, C. M., de Almeida Mantovani, D. B., & da Silva, A. M. M. (2020). Resting state brain in cognitive decline: an analysis of brain network architecture using graph theory. Anais do CBEB2020, 2020, Brasil..

Newman, M. E. (2004). Fast algorithm for detecting community structure in networks. Physical review E, 69 (6), 066133.

Power, J. D., Barnes, K. A., Snyder, A. Z., Schlaggar, B. L., & Petersen, S. E. (2012). Spurious but systematic correlations in functional connectivity mri networks arise from subject motion. Neuroimage, 59 (3), 2142–2154.

Power, J. D., Mitra, A., Laumann, T. O., Snyder, A. Z., Schlaggar, B. L., & Petersen, S. E. (2014). Methods to detect, characterize, and remove motion artifact in resting state fmri. Neuroimage, 84, 320–341.

Ran, Q., Jamoulle, T., Schaeverbeke, J., Meersmans, K., Vandenberghe, R., & Dupont, P. (2020). Reproducibility of graph measures at the subject level using resting-state fmri. Brain and behavior, 10 (8), 2336–2351.

Rubinov, M., & Sporns, O. (2010). Complex network measures of brain connectivity: uses and interpretations. Neuroimage, 52 (3), 1059–1069.

Sanz-Arigita, E. J., Schoonheim, M. M., Damoiseaux, J. S., Rombouts, S. A., Maris, E., Barkhof, F., … Stam, C. J. (2010). Loss of ‘small-world’networks in alzheimer’s disease: graph analysis of fmri resting-state functional connectivity. PloS one, 5 (11), e13788.

Schwarz, A. J., & McGonigle, J. (2011). Negative edges and soft thresholding in complex network analysis of resting state functional connectivity data. Neuroimage, 55 (3), 1132–1146.

Shirer, W. R., Ryali, S., Rykhlevskaia, E., Menon, V., & Greicius, M. D. (2012). Decoding subject-driven cognitive states with whole-brain connectivity patterns. Cerebral cortex, 22 (1), 158–165.

Si, S.-Z., Liu, X., Wang, J.-F., Wang, B., & Zhao, H. (2019). Brain networks modeling for studying the mechanism underlying the development of alzheimer’s disease. Neural regeneration research, 14 (10), 1805.

Sperling, R. A., Aisen, P. S., Beckett, L. A., Bennett, D. A., Craft, S., Fagan, A. M., … others (2011). Toward defining the preclinical stages of alzheimer’s disease: Recommendations from the national institute on aging-alzheimer’s association workgroups on diagnostic guidelines for alzheimer’s disease. Alzheimer’s & dementia, 7 (3), 280–292.

Subramanian, S., Rajamanickam, K., Prakash, J. S., Ramachandran, M., (Adni, A. D. N. I., et al. (2020). Study on structural atrophy changes and functional connectivity measures in alzheimer’s disease. Journal of Medical Imaging, 7 (1), 016002.

Supekar, K., Menon, V., Rubin, D., Musen, M., & Greicius, M. D. (2008). Network analysis of intrinsic functional brain connectivity in alzheimer’s disease. PLoS computational biology, 4 (6), e1000100.

Van Dijk, K. R., Sabuncu, M. R., & Buckner, R. L. (2012). The influence of head motion on intrinsic functional connectivity mri. Neuroimage, 59 (1), 431–438.

Vy’tvarová, E., Fousek, J., Bartoň, M., Mareček, R., Gajdoš, M., Lamoš, M., … Mikl, M. (2017). The impact of diverse preprocessing pipelines on brain functional connectivity. In 2017 25th european signal processing conference (eusipco) (pp. 2644–2648).

Wang, J., Wang, L., Zang, Y., Yang, H., Tang, H., Gong, Q., … He, Y. (2009). Parcellation-dependent small-world brain functional networks: A resting-state fmri study. Human brain mapping, 30 (5), 1511–1523.

Wang, J., Zuo, X., Dai, Z., Xia, M., Zhao, Z., Zhao, X., … He, Y. (2013). Disrupted functional brain connectome in individuals at risk for alzheimer’s disease. Biological psychiatry, 73 (5), 472–481.

Watts, D. J., & Strogatz, S. H. (1998). Collective dynamics of ‘small-world’networks. nature, 393 (6684), 440–442.

Weis, C. N., Bennett, K. P., Huggins, A. A., Parisi, E. A., Gorka, S. M., & Larson, C. (2021). A 7-tesla mri study of the periaqueductal gray: resting state and task activation under threat. Social Cognitive and Affective Neuroscience.

Xiang, J., Guo, H., Cao, R., Liang, H., & Chen, J. (2013). An abnormal resting-state functional brain network indicates progression towards alzheimer’s disease. Neural regeneration research, 8 (30), 2789.

Xu, X., Li, W., Mei, J., Tao, M., Wang, X., Zhao, Q., … Wang, P. (2020). Feature selection and combination of information in the functional brain connectome for discrimination of mild cognitive impairment and analyses of altered brain patterns. Frontiers in aging neuroscience, 12, 28.

Xue, C., Sun, H., Hu, G., Qi, W., Yue, Y., Rao, J., … others (2020). Disrupted patterns of rich-club and diverse-club organizations in subjective cognitive decline and amnestic mild cognitive impairment. Frontiers in neuroscience, 984.

Yan, C.-G., Craddock, R. C., He, Y., & Milham, M. P. (2013). Addressing head motion dependencies for small-world topologies in functional connectomics. Frontiers in human neuroscience, 7, 910.

Zhang, C., Cahill, N. D., Arbabshirani, M. R., White, T., Baum, S. A., & Michael, A. M. (2016). Sex and age effects of functional connectivity in early adulthood. Brain connectivity, 6 (9), 700–713.

Zhang, L., Ni, H., Yu, Z., Wang, J., Qin, J., Hou, F., … others (2020). Investigation on the alteration of brain functional network and its role in the identification of mild cognitive impairment. Frontiers in neuroscience, 14, 1027.

Zhao, X., Liu, Y., Wang, X., Liu, B., Xi, Q., Guo, Q., … Wang, P. (2012). Disrupted small-world brain networks in moderate alzheimer’s disease: a resting-state fmri study. PloS one, 7 (3), e33540.

