## Supplemental Material for "Denoising Approach Affects Diagnostic Differences in Brain Connectivity across the Alzheimer’s Disease Continuum"

### Supplementary Material

#### Table 1

***Supplemental Table 1**. ADNI Participant and Scan IDs.*

| RID | Image ID | $RID_{cont.}$ | Image $ID_{cont.}$ | $RID_{cont.}$ | Image $ID_{cont.}$ | $RID_{cont.}$ | Image $ID_{cont.}$ |
|---|---|---|---|---|---|---|---|
| 413 | 863058 | 4597 | 290305 | 4464 | 959744 | 4891 | 1027772 |
| 4225 | 1075138 | 4696 | 301395 | 6200 | 964919 | 259 | 907415 |
| 4654 | 848004 | 4868 | 319541 | 6306 | 988750 | 1427 | 901033 |
| 5178 | 857837 | 4252 | 258955 | 6480 | 1017995 | 2234 | 955111 |
| 6103 | 938771 | 4367 | 269279 | 6551 | 1037230 | 5185 | 890736 |
| 6456 | 1017840 | 4549 | 300334 | 377 | 1117318 | 5200 | 871946 |
| 1074 | 912462 | 4680 | 300529 | 454 | 900940 | 2002 | 1221691 |
| 4119 | 1249340 | 4835 | 315850 | 4302 | 812267 | 2130 | 1049546 |
| 4441 | 1154574 | 5012 | 343912 | 4308 | 894408 | 778 | 334140 |
| 6256 | 973282 | 6185 | 958015 | 4410 | 824984 | 2332 | 852334 |
| 6258 | 973297 | 626 | 886319 | 4510 | 914180 | 4287 | 343366 |
| 6260 | 1001100 | 4276 | 886317 | 4513 | 931964 | 4422 | 317121 |
| 6264 | 1005760 | 4659 | 887787 | 4974 | 915136 | 6763 | 1186907 |
| 6268 | 1005896 | 5236 | 886309 | 5026 | 401558 | 6852 | 1285630 |
| 6479 | 1246024 | 5237 | 1036198 | 5078 | 882556 | 2373 | 302589 |
| 6644 | 1083055 | 6312 | 988050 | 5097 | 971714 | 2403 | 341127 |
| 6833 | 1254311 | 4674 | 880429 | 6731 | 1177872 | 4294 | 267894 |
| 4150 | 249407 | 120 | 838962 | 6785 | 1219611 | 4417 | 279181 |
| 4153 | 248516 | 2219 | 840547 | 6801 | 1227910 | 4542 | 287650 |
| 4192 | 258605 | 5118 | 875808 | 4813 | 316519 | 4660 | 300043 |
| 4346 | 266131 | 4290 | 1133906 | 5070 | 357475 | 4730 | 306375 |

| RID | Image ID | $RID_{cont.}$ | Image $ID_{cont.}$ | $RID_{cont.}$ | Image $ID_{cont.}$ | $RID_{cont.}$ | Image $ID_{cont.}$ |
|---|---|---|---|---|---|---|---|
| 4357 | 268917 | 4384 | 990074 | 934 | 1033065 | 4817 | 314327 |
| 4363 | 269256 | 5219 | 1146498 | 5292 | 1047959 | 4925 | 337131 |
| 4713 | 303733 | 2233 | 275532 | 210 | 939839 | 4971 | 342326 |
| 4867 | 322009 | 4021 | 246871 | 2184 | 920084 | 4982 | 341793 |
| 6303 | 985201 | 4024 | 228872 | 2187 | 1036868 | 4984 | 342278 |
| 6465 | 1041486 | 4042 | 235238 | 4067 | 892725 | 4990 | 342915 |
| 4012 | 306576 | 4203 | 255309 | 4332 | 874429 | 4997 | 347402 |
| 4128 | 265533 | 4218 | 255986 | 4431 | 913458 | 5006 | 348491 |
| 4188 | 255318 | 4590 | 290644 | 2121 | 963758 | 4489 | 858987 |
| 4545 | 312870 | 4721 | 304675 | 4428 | 947591 | 4189 | 330141 |
| 6073 | 908590 | 4947 | 339436 | 5282 | 1004666 | 2333 | 949423 |
| 4580 | 296863 | 5289 | 849894 | 6287 | 998051 | 6062 | 892773 |
| 4595 | 300057 | 6279 | 981034 | 6690 | 1162193 | 6151 | 947548 |
| 5071 | 360323 | 6293 | 993281 | 4003 | 943601 | 6426 | 1015402 |
| 5137 | 368206 | 6700 | 1154756 | 6632 | 1091695 | 6467 | 1017242 |
| 5171 | 374972 | 6804 | 1230903 | 4404 | 949880 | 6821 | 1244554 |
| 2133 | 280337 | 4176 | 1117702 | 6347 | 1011831 | 6828 | 1240486 |
| 2155 | 274112 | 555 | 964901 | 4043 | 969405 | 6843 | 1275419 |
| 4349 | 266634 | 4414 | 915043 | 4507 | 1003343 | 4100 | 923855 |

**Table 2**

***Supplemental Table 2***. *ROI Anatomical Information.*

| Networks | N Voxels | Anatomical Region |
|---|---|---|
| PRE | 165 | Cingulate Cortex (Middle/Posterior) |
| | 469 | Precuneus |
| | 118 | Left Inferior/Superior Parietal Lobule |
| | 30 | Right Inferior/Superior Parietal Lobule |
| VDMN | 134 | Left PCC/Precuneus |
| | 122 | Left Middle/Superior Frontal Gyrus |
| | 40 | Left Fusiform/Parahippocampal Gyrus |
| | 149 | Left Superior Occipital Gyrus |
| | 176 | Right PCC |
| | 574 | Precuneus |
| | 115 | Right Middle/Superior Frontal Gyrus |
| | 24 | Right Cerebellum (IV-V) |
| | 221 | Right Middle Temporal Gyrus |
| DDMN | 1551 | Left Medial/Superior Frontal Gyrus |
| | 31 | Left Angular Gyrus |
| | 43 | Right Superior Frontal Gyrus |
| | 460 | Precuneus |
| | 36 | Middle Cingulate Cortex |
| | 11 | Right Angular Gyrus |
| | 58 | Thalamus-Proper |
| | 120 | Left Parahippocampal Gyrus |
| | 38 | Right Parahippocampal Gyrus |
| LECN | 441 | Left Superior/Middle Frontal Gyrus |

| Networks | N Voxels | Anatomical Region |
|---|---|---|
| | 121 | Left Middle/Inferior Frontal Gyrus |
| | 621 | Left Inferior Parietal Lobule |
| | 3 | Left Thalamus |
| RECN | 619 | Right Middle Frontal Gyrus |
| | 106 | Right Middle Frontal Gyrus |
| | 546 | Right Inferior Parietal Lobule |
| | 22 | Right Superior/Medial Frontal Gyrus |
| | 711 | Left Cerebellum (Crus II/I) |
| | 53 | Right Caudate Nucleus |
| VSPN | 101 | Left Middle Frontal Gyrus |
| | 588 | Left Inferior/Superior Parietal Lobule |
| | 329 | Left Inferior/Middle Frontal Gyrus |
| | 32 | Left Middle Occipital Gyrus |
| | 30 | Right Middle Frontal Gyrus |
| | 360 | Right Superior/Inferior Parietal Lobule |
| | 92 | Right Inferior/Middle Frontal Gyrus |
| | 22 | Right Fusiform Gyrus |
| | 19 | Right Cerebellum (Crus I) |
| ASAL | 204 | Left Middle/Superior Frontal Gyrus |
| | 88 | Left Insula |
| | 834 | Left/Right Superior Frontal Gyrus |
| | 144 | Right Superior/Middle Frontal Gyrus |
| | 97 | Right Insula/Inferior Frontal Gyrus |
| | 38 | Right Cerebellum (Crus I) |
| PSAL | 26 | Left Middle Frontal Gyrus |

| Networks | N Voxels | Anatomical Region |
|---|---|---|
| | 354 | Left Inferior Parietal Lobule |
| | 28 | Left Precuneus |
| | 14 | Right Paracentral Lobule |
| | 44 | Right Postcentral Gyrus |
| | 297 | Right Inferior Parietal Lobule |
| | 46 | Left Thalamus |
| | 31 | Left Insula |
| | 22 | Right Thalamus |
| | 41 | Right Insula |
| LAN | 192 | Left Inferior Frontal Gyrus |
| | 102 | Left Middle Temporal Gyrus |
| | 419 | Left Superior/Middle Temporal Gyrus |
| | 15 | Right Inferior Frontal Gyrus |
| | 337 | Right Superior/Middle Temporal Gyrus |
| SMN | 417 | Left Postcentral/Precentral Gyrus |
| | 414 | Right Pre/Postcentral Gyrus |
| | 42 | Right Medial Frontal Gyrus |
| | 4 | Left Thalamus |
| | 594 | Left/Right Cerebellum (VI) / Cerebellar Vermis |
| | 5 | Right Thalamus |
| PVIS | 327 | Left/Right Cuneus |
| | 1 | Thalamus |
| HVIS | 251 | Left Middle/Inferior Occipital Lobe |
| | 498 | Right Middle/Inferior Occipital Gyrus |
| AUD | 284 | Left Superior Temporal Gyrus |

| Networks | N Voxels | Anatomical Region |
|---|---|---|
| | 160 | Right Superior Temporal Gyrus |
| | 8 | Right Thalamus |
| BG | 192 | Right Thalamus |
| | 237 | Left Thalamus |
| | 7 | Left Middle Frontal Gyrus |
| | 19 | Right Middle/Inferior Frontal Gyrus |

PRE = precuneus network; VDMN = ventral default mode network; DDMN = dorsal default mode network; LECN = left executive control network; RECN = right executive control network; VSPN = visuospatial network; ASAL = anterior salience network; PSAL = posterior salience network; LAN = language network; SMN = sensorimotor network; PVIS = primary visual network; HVIS = higher visual network; AUD = auditory network; BGN = basal ganglia network.
